# KSHV miR-K12-9 Induces Transformation of Immortalized and Primary Endothelial Cells

**DOI:** 10.64898/2026.05.18.726106

**Authors:** Lauren A. Gay, Vijay Sirohi, Melody Baddoo, Erik Flemington, Scott Tibbetts, Rolf Renne

## Abstract

Like most herpesviruses, KSHV encodes multiple microRNAs (miRNAs). Collectively, they comprise an important mechanism through which the virus maintains latency and persists in cells. At the same time, individual miRNAs can also play distinct, nonredundant roles. Past experiments with single miRNA knockout viruses showed that miR-K12-9, in particular, filled a unique niche. Endothelial cells latently infected with the miR-K12-9 knockout grew to be many times larger than WT-infected cells and proliferated at a significantly slower rate. Their ability to migrate was slowed as well. RNA-seq identified nearly 8,500 differentially expressed genes between miR-K12-9 knockout- and WT-infected cells. To further study miR-K12-9, we generated Telomerase-Immortalized Microvascular Endothelial (TIME) cells expressing either miR-K12-9 or a control miRNA from a lentivirus. Unexpectedly, after approximately one month in culture, unmistakable morphological changes began to occur in two of the three miR-K12-9-expressing cell lines. These smaller, more rounded cells proliferated rapidly and swiftly took over the two cultures. Given this result, we proceeded to characterize all the lentivirus-transduced cell lines in various assays focused on oncogenesis. When looking at colony formation in soft agar, only those two miR-K12-9-expressing cell lines produced colonies, indicating a loss of contact inhibition. NOD/SCID mice injected with the two cell miR-K12-9-expressing cell lines developed tumors while those receiving other cell lines did not. To confirm reproducibility of these results, we transduced both TIME and primary endothelial cells (HUVECs) with the miR-K12-9 and control lentiviruses. Once again, approximately half of the cell lines expressing miR-K12-9 showed hallmark phenotypes of transformation. We are currently characterizing the miR-K12-9 targetome in the transduced cell lines and mouse tumors using bulk and single-cell RNA-seq. This should yield insights into the underlying mechanism and required cofactors of miR-K12-9-induced transformation. To our knowledge, this is the first description of transformation of endothelial cells by a viral miRNA.

## Introduction

An oncogenic virus, Kaposi’s Sarcoma-Associated Herpesvirus (KSHV) is the etiological agent of Kaposi’s Sarcoma (KS), Primary Effusion Lymphoma (PEL), some forms of Multicentric Castleman’s Disease (MCD), and KSHV inflammatory cytokine syndrome (KICS) (1,2,3,4). KSHV is a gammaherpesvirus with a genome of approximately 140bp that encodes both protein and noncoding RNA. Like other herpesviruses, KSHV undergoes lytic and latent replication cycles characterized by distinct genetic programs. The majority of genes are expressed during the lytic phase, in which there is active production of viral progeny. By contrast, latency is a quiescent state in which the viral genome passively replicates with each cell division (5). This phase is associated with oncogenesis (6). Among the small group of genes expressed during latency are twelve miRNAs (7,8,9).

MiRNAs are a class of 19-22bp long RNAs that regulate cellular processes by repressing target genes. They originate in the nucleus as primary transcripts (pri-miRNAs) containing one or more hairpin, or stem-loop, structures. These are recognized and trimmed by Drosha/DGCR8 into single hairpin pre-miRNAs. The pre-miRNAs are exported into the cytoplasm where they encounter Dicer, an endonuclease which cleaves the loop from the pre-miRNA, releasing a short RNA duplex consisting of the 5’ and 3’ arms of the hairpin. One of the arms becomes incorporated into the RNA-Induced Silencing Complex (RISC) while the other is released. This process is often asymmetrical, with one strand preferred over the other. Once the RISC is assembled, the miRNA binds to partially complementary sequences in target transcripts. This results in translational inhibition and/or degradation of the target RNA (10).

MiRNAs are known to bind both coding and noncoding transcripts (11,12). The function of a miRNA is defined by the genes it targets for downregulation. Multiple KSHV miRNAs have been found to play a role in oncogenesis through targeting genes and pathways regulating this process (13). One salient example is miR-K12-11, which shares a seed sequence with a cellular pro-oncogenic miRNA, miR-155 (14,15). Ectopic expression of either miR-K12-11 or miR-155 in splenic B-cells or bone marrow-derived hematopoietic stem cells resulted in significant expansions of these cell populations (16,17). It has been reported that miR-K12-3 directly targets G protein-coupled receptor kinase 2 (GRK2) to promote migration and invasion of endothelial cells (18). KSHV-infected cells show resistance to apoptosis, at least due in part to the targeting of Caspase-3 by miR-K12-1, -3, and –4-3p (19). Due to their short recognition sequence, miRNAs never target just one gene. Instead, they serve as node points in a regulatory web.

In earlier work, we described the oncogenesis-related phenotypic characteristics of endothelial cells infected with a panel of miRNA knockout viruses. We observed that individual knockout-infected cell lines showed unique phenotypic traits that allowed them to be distinguished from one another. This suggests that each viral miRNA has its own specific contribution to the KSHV replication cycle and virally induced oncogenesis. At the time, it was noted that the absence of one miRNA, miR-K12-9, had particularly profound effects on infected cells. The miR-K12-9 KO mutant significantly decreased the proliferation rate while dramatically increasing cell size (19).

Although not previously implicated in oncogenesis directly, miR-K12-9 is known to contribute to immune evasion, protection from apoptosis, and the maintenance of latency. MiR-K12-9-3p targets IRAK1 (interleukin 1 receptor-associated kinase 1) to interfere with the activation of NF-KB and MAP kinases, reducing the downstream expression of pro-inflammatory cytokines such as IL-6 (20). It also directly downregulates GADD45B (growth arrest DNA damage-inducible gene 45 beta), preventing cell cycle arrest and apoptosis (21). At the same time, miR-K12-9-5p, along with miR-K12-7-5p, targets RTA, the viral reactivation switch, leading to the suppression of lytic replication (22,23). MiR-K12-9 has a high degree of polymorphism, with SNPs identified in the pri-, pre- and mature miRNA forms (24,25,26). Variations in miR-K12-9 and throughout the miRNA locus are associated with lower expression of miR-K12-9 (26).

High levels of polymorphism in the miRNA region are characteristic of KSHV isolates from KSHV-MCD and KICS patients (24,27). Unlike KS, KSHV-MCD and KICS show extensive lytic replication and strong immune activation characterized by the overproduction of IL-6 (4,28). The principal features of these two diseases may relate to low expression of miR-K12-9 and the subsequent loss of control over lytic replication and the innate immune response.

Here we present RNA-seq and small RNA-seq datasets from the complete panel of KSHV miRNA KO mutants in endothelial cells, a relevant model for KS. Like the mutant viruses themselves, these data are intended to serve as a resource for the broader KSHV research community. All sequencing data can be accessed on GEO. Furthermore, the unusual characteristics of miR-K12-9 KO-infected cells prompted us to continue our line of inquiry concerning this miRNA. Unexpectedly, the sustained ectopic expression of miR-K12-9 in either immortalized or primary endothelial cells resulted in transformation 50% of the time. To the best of our knowledge, this is the first report of a KSHV miRNA capable of driving oncogenesis in endothelial cells.

## Results

Previously, we infected telomerase-immortalized vein endothelial (TIVE) cells with a panel of miRNA knockout mutants. Once latency was established, the mutant cell lines were compared to WT-infected TIVE cells in phenotypic assays related to oncogenesis. In the present work we made use of these already established cell lines and performed RNA-seq and small RNA-seq. Differential expression analysis comparing WT-infected cells with each miRNA KO-infected cell line presented an opportunity to further tease apart the specific role of each KSHV miRNA in the viral replication cycle and oncogenesis. The results indicate that, as a group, the KSHV miRNAs are involved in diverse cellular pathways, while on an individual level, they exhibit both redundancy and specialization. Given the size and complexity of the full dataset from the complete panel of miRNA knockout mutants, only a few of the miRNAs are highlighted here. Additional results are provided in supplementary figures S1-S7. In parallel, small RNA sequencing was performed on the same cell lines to assess the impact of KSHV miRNA deletion on host miRNA expression. While detailed analysis was not pursued here, global patterns suggested that loss of individual viral miRNAs was associated with modest but detectable changes in host miRNA profiles (Fig. S8).

Prior work demonstrated that knockout of either miR-K12-5 and miR-K12-7 results in a pronounced impairment in tubule formation, a commonly used proxy for angiogenic capacity, suggesting that both of these miRNAs are particularly important for sustaining this process. Angiogenesis is critically dependent on dynamic interactions between endothelial cells and the surrounding extracellular matrix (ECM), which provides both structural support and biochemical cues necessary for endothelial migration, signaling, and structural organization (29). Consistent with this functional phenotype, differential expression analysis revealed distinct but selective transcriptional changes in both miR-K12-5 and miR-K12-7 knockout cells relative to WT controls (Fig. 1A, B). To determine whether these gene-level changes converged on specific biological processes, gene set enrichment analysis was performed (Fig. 1D, E). This identified strong enrichment of pathways associated with the ECM, including extracellular structure organization, external encapsulating structure organization, and ECM organization. It may seem paradoxical that the loss of two seemingly pro-angiogenic miRNAs would be associated with upregulation, rather than downregulation, of collagens and other ECM-related genes. However, the ECM plays a complex and dynamic role in angiogenesis, acting not only as a structural scaffold but also as a source of regulatory signals that can either promote or inhibit vascular growth depending on its composition and remodeling state (30). Indeed, collagens regulate angiogenesis through both intact structural functions and proteolytically generated fragments with distinct pro- or anti-angiogenic activities (31).

**Figure 1.**
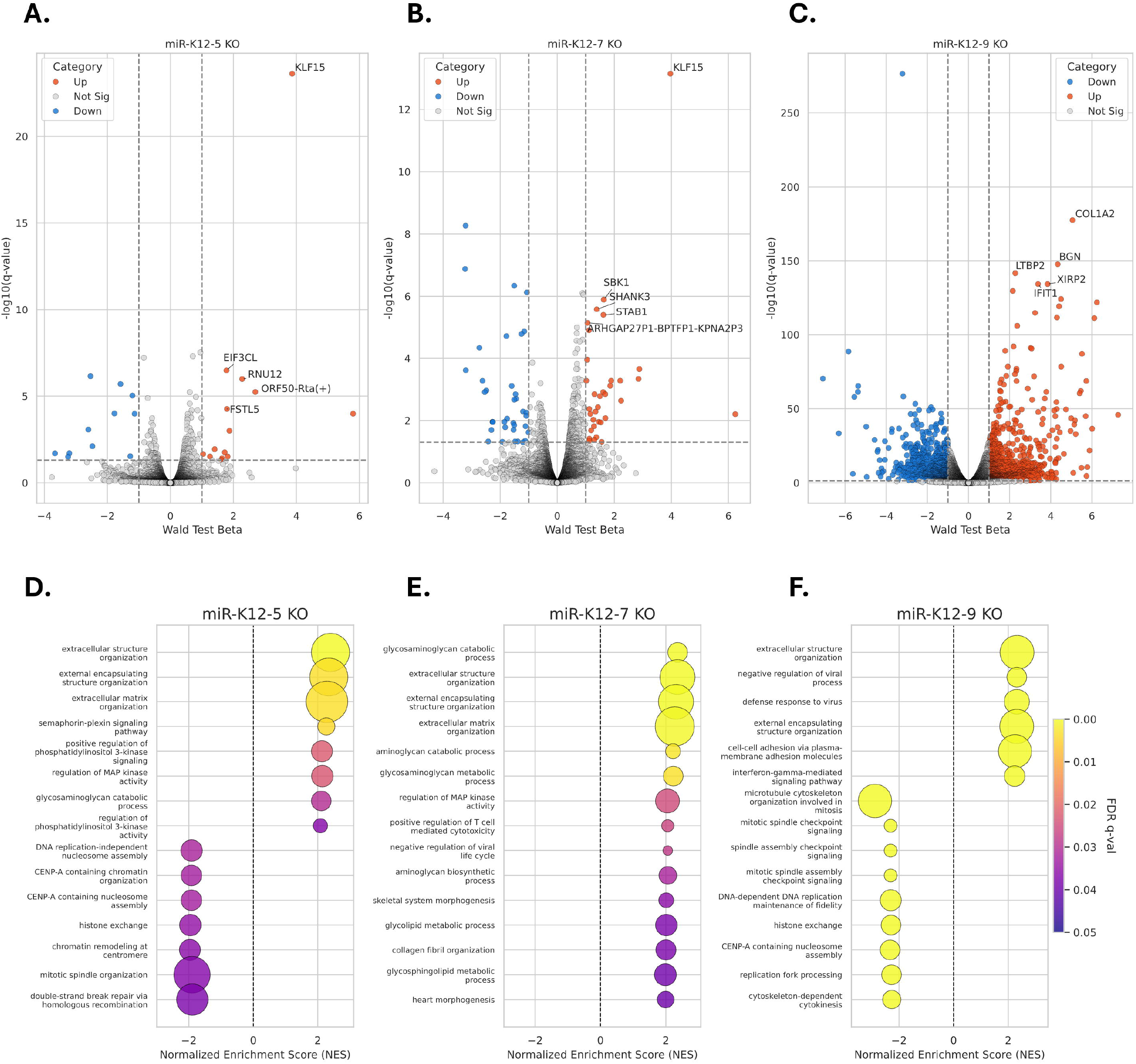
Representative RNA-seq analysis results showing the miR-K12-5, -7, and -9 KO mutants vs. WT. A. Differential expression analysis was performed with Sleuth. Volcano plots show the magnitude of expression change (Wald test beta) vs. significance. Genes in blue are significantly downregulated, genes in red are significantly upregulated, and genes in gray have no significant change in expression. Top 5 upregulated genes are labeled. B. Bubble plots show the results of gene set enrichment analysis. Negative enrichment scores indicate that genes in the set are downregulated while positive enrichment scores indicate the opposite. The size of each bubble corresponds to the number of genes in the set while the color indicates the level of significance. RNA-seq was performed in triplicate using RNA from 3 separate preparations.

In contrast to the more selective transcriptional changes observed for other miRNA knockouts, the miR-K12-9 knockout exhibited a markedly divergent gene expression profile relative to WT-infected cells. Differential expression analysis identified over 8,000 significantly altered genes (Fig. 1C; Fig. 2), exceeding the combined total observed across the other miRNA mutants and indicating a broad, global disruption of host transcriptional programs. Consistent with this level of change, gene set enrichment analysis revealed enrichment across a wide range of biological pathways (Fig. 1F), including extracellular matrix organization, immune and antiviral signaling, and cell cycle-associated processes. Notably, while ECM-related pathways were also enriched in other knockouts, the miR-K12-9 mutant was distinguished by the simultaneous and pronounced involvement of inflammatory and proliferative signaling networks. This broad and multifaceted transcriptional reprogramming underscores the unique role of miR-K12-9 among KSHV miRNAs, consistent with its distinct phenotypic effects observed in prior studies, and suggests that miR-K12-9 may function as a key regulator of global cellular state rather than modulating a more restricted set of pathways.

**Figure 2.**
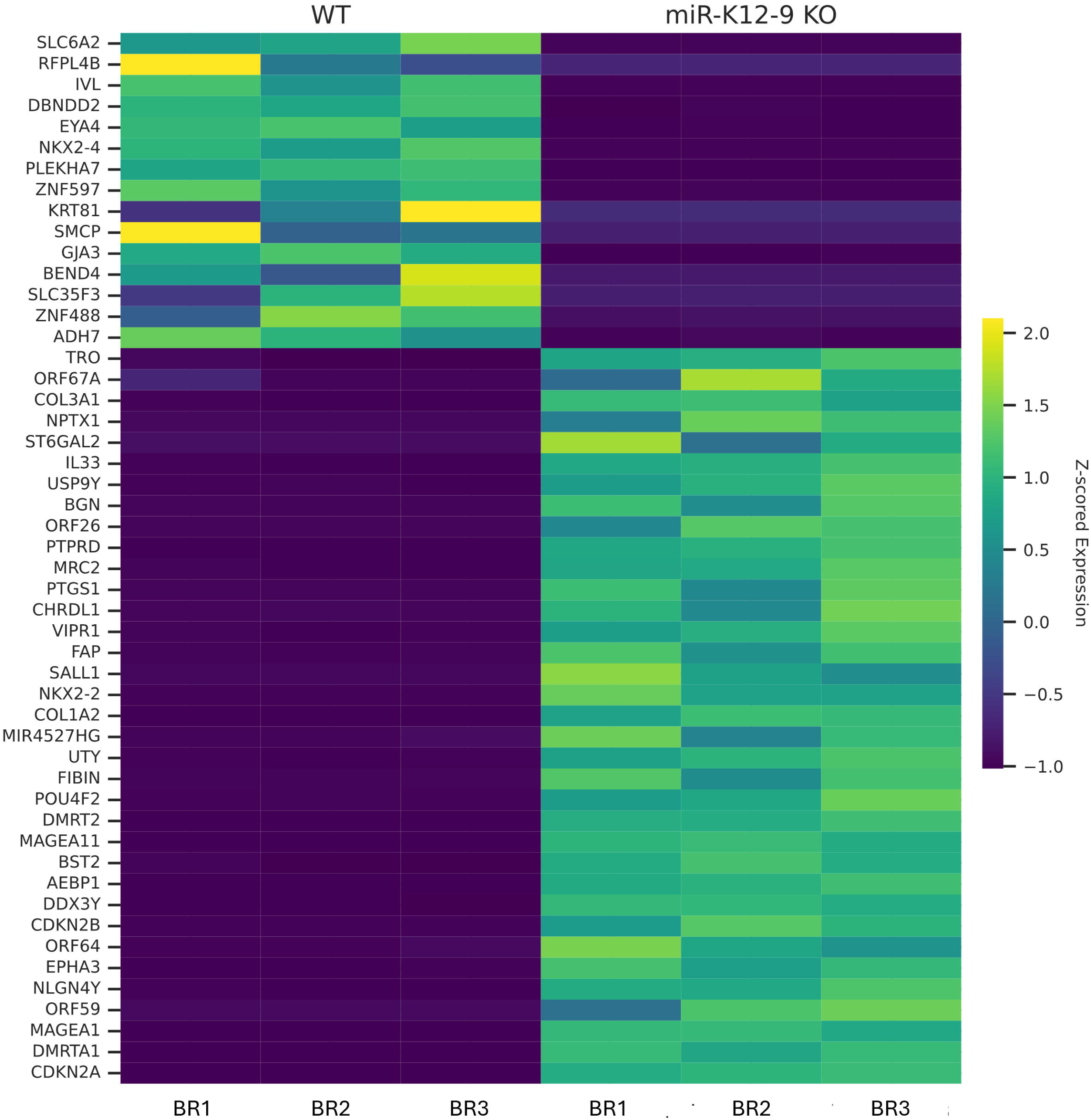
Heatmap showing the top 50 differentially expressed genes between the miR-K12-9 KO mutant and WT.

Given the unusually broad transcriptional effects observed following miR-K12-9 knockout, coupled with previous observations, we chose to focus subsequent studies on this miRNA. However, technical challenges limited further use of the original knockout cell line, necessitating an alternative approach. To directly assess the functional impact of miR-K12-9, we generated a lentiviral construct expressing full-length miR-K12-9 under the control of the U6 promoter, along with a scrambled miRNA control. Telomerase-immortalized microvascular endothelial (TIME) cells were transduced in triplicate and selected to establish stable cell populations.

During expansion of the transduced cells, distinct morphological changes were observed in a subset of the miR-K12-9-expressing cultures. Specifically, dense clusters of small, rapidly proliferating cells emerged and progressively overtook the surrounding monolayer (Fig. 3A). This phenotype was reproducible across independent experiments, arising in multiple biological replicates (Fig. 3B), suggesting that miR-K12-9 expression can drive a stable and heritable alteration in cell growth behavior. Expression of mature miR-K12-9-5p and -3p was confirmed by qPCR (Fig. 4A, 4B), although absolute expression levels did not strictly correlate with the emergence of the phenotype.

**Figure 3.**
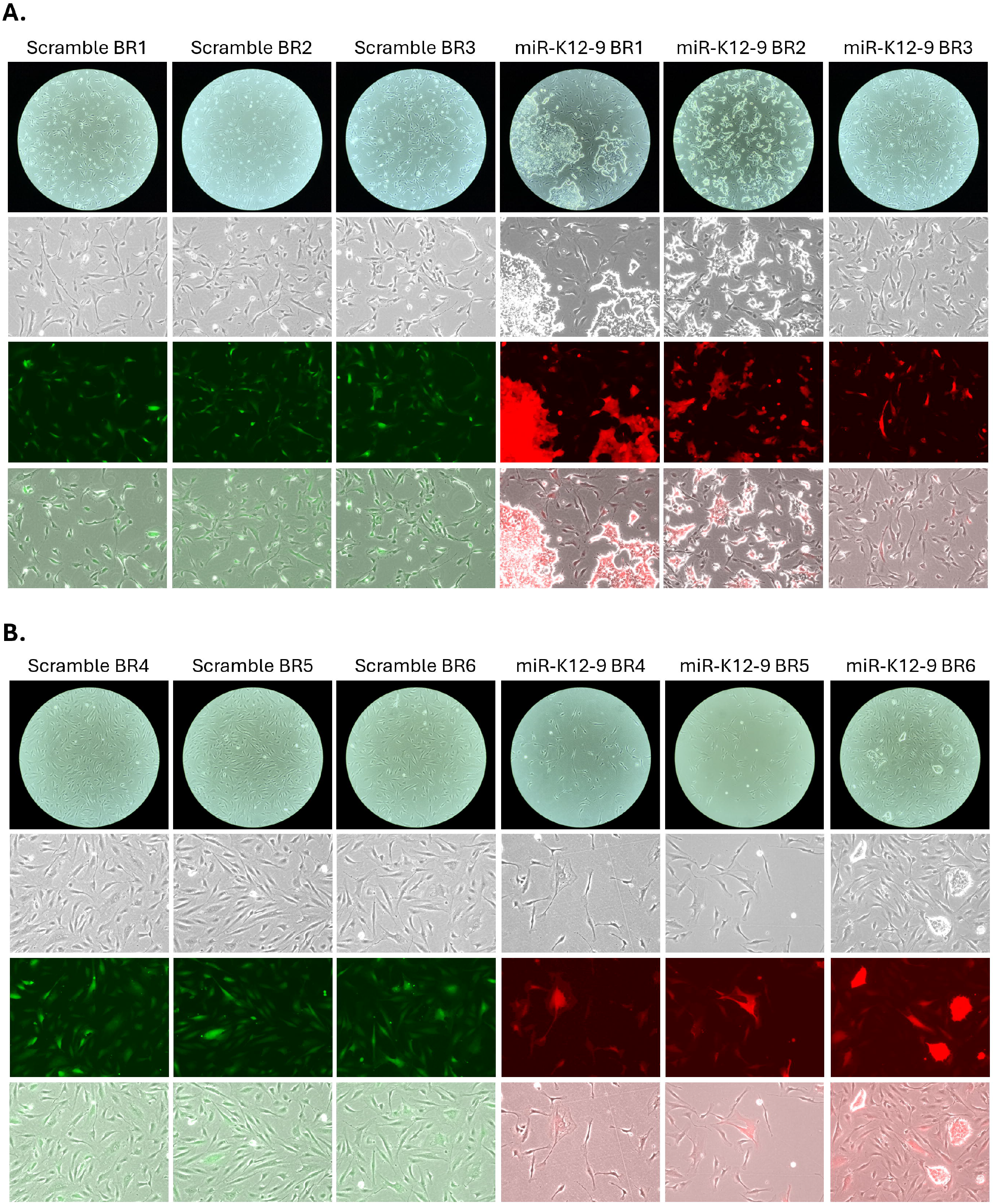
Heatmap showing the top 50 differentially expressed host miRNAs across all KSHV miRNA KO mutants.

**Figure 4.**
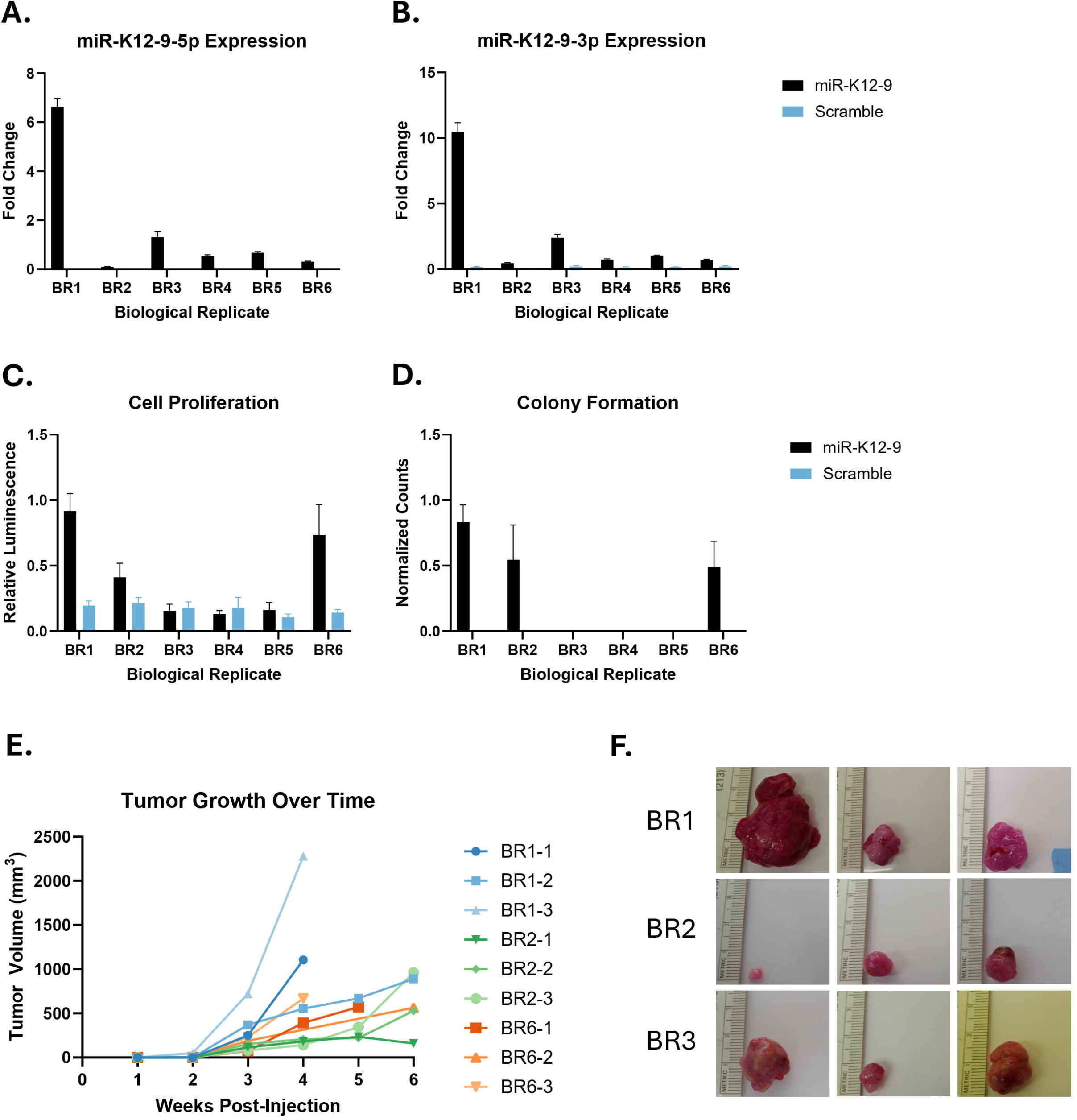
Images of TIME cells transduced with control (GFP) and miR-K12-9 (mCherry) lentiviruses. TIME cells in 6-well plates were transduced in triplicate with either control miRNA- or miR-K12-9-expressing lentivirus. Each transduced well is considered a biological replicate (BR). A. Photographs of the first experiment, taken 39 days post-transduction, after puromycin selection and expansion into 10cm plates. B. Photographs of the second experiment, taken 23 days post-transduction, after puromycin selection and expansion into 10cm plates.

To further characterize this observation, we assessed hallmark features of cellular transformation. In proliferation assays, a subset of miR-K12-9-expressing lines (BR1, BR2, and BR6) displayed significantly increased growth rates relative to both control cells and miR-K12-9 cells without phenotypic changes (Fig. 4C). Consistent with this, these same replicates demonstrated robust colony formation in soft agar, indicative of anchorage-independent growth, whereas control cells and the remaining miR-K12-9-expressing lines failed to form colonies (Fig. 4D).

To determine whether these in vitro phenotypes extended to tumorigenic potential in vivo, cells were evaluated in a xenograft model using NOD/SCID mice. Notably, only mice injected with miR-K12-9-expressing BR1, BR2, and BR6 developed tumors, while control cells and the remaining replicates did not produce detectable growths. Tumor growth kinetics further distinguished these lines, with BR1 exhibiting the most aggressive growth, followed by BR6 and BR2. In some cases, tumor burden necessitated early sacrifice due to size limits (Fig. 4E, F).

To further examine the transcriptional composition of miR-K12-9-driven tumors, single-cell RNA sequencing was performed on tumors derived from BR1, BR2, and BR6 cell lines, henceforth referred to as the transformed cell lines. Unsupervised clustering revealed a high degree of similarity across biological replicates, with no clear separation based on the originating cell line. However, tumor cells consistently segregated into a small number of transcriptionally distinct clusters, suggesting the presence of intratumoral heterogeneity. The extent of transcriptional divergence between these clusters appeared modest, and in the absence of single-cell profiling of the parental cell populations, it remains unclear whether this heterogeneity reflects pre-existing cellular diversity or phenotypic diversification following tumor formation. These findings indicate that miR-K12-9-driven tumors adopt a broadly consistent transcriptional state while retaining limited cellular heterogeneity.

Together, these findings demonstrate that ectopic expression of miR-K12-9 is sufficient to drive cellular transformation in a subset of endothelial cell populations, as evidenced by increased proliferation, anchorage-independent growth, and tumor formation in vivo. Notably, while the presence of the transformed phenotype did not strictly correlate with miR-K12-9 expression levels, the magnitude of the phenotype was proportional to expression, with higher-expressing lines consistently exhibiting more aggressive behavior. These results identify miR-K12-9 as a potent driver of oncogenic transformation and suggest that, in the context of KSHV infection, this miRNA may play a central role in promoting tumorigenesis in endothelial cells.

To further define the transcriptional consequences of miR-K12-9 expression, RNA-seq was performed on all lentivirus-transduced TIME cell lines. Unsupervised principal component analysis revealed clear segregation of samples based on both miR-K12-9 expression and transformation status (Figure 5A). Control (scramble) samples clustered tightly, indicating a consistent baseline transcriptional state, whereas miR-K12-9-expressing cells were distinctly separated along the primary axis of variation. Notably, non-transformed miR-K12-9-expressing cells occupied an intermediate position between control and transformed samples, while transformed cells formed a more distant and cohesive cluster, consistent with a more pronounced transcriptional shift.

**Figure 5.**
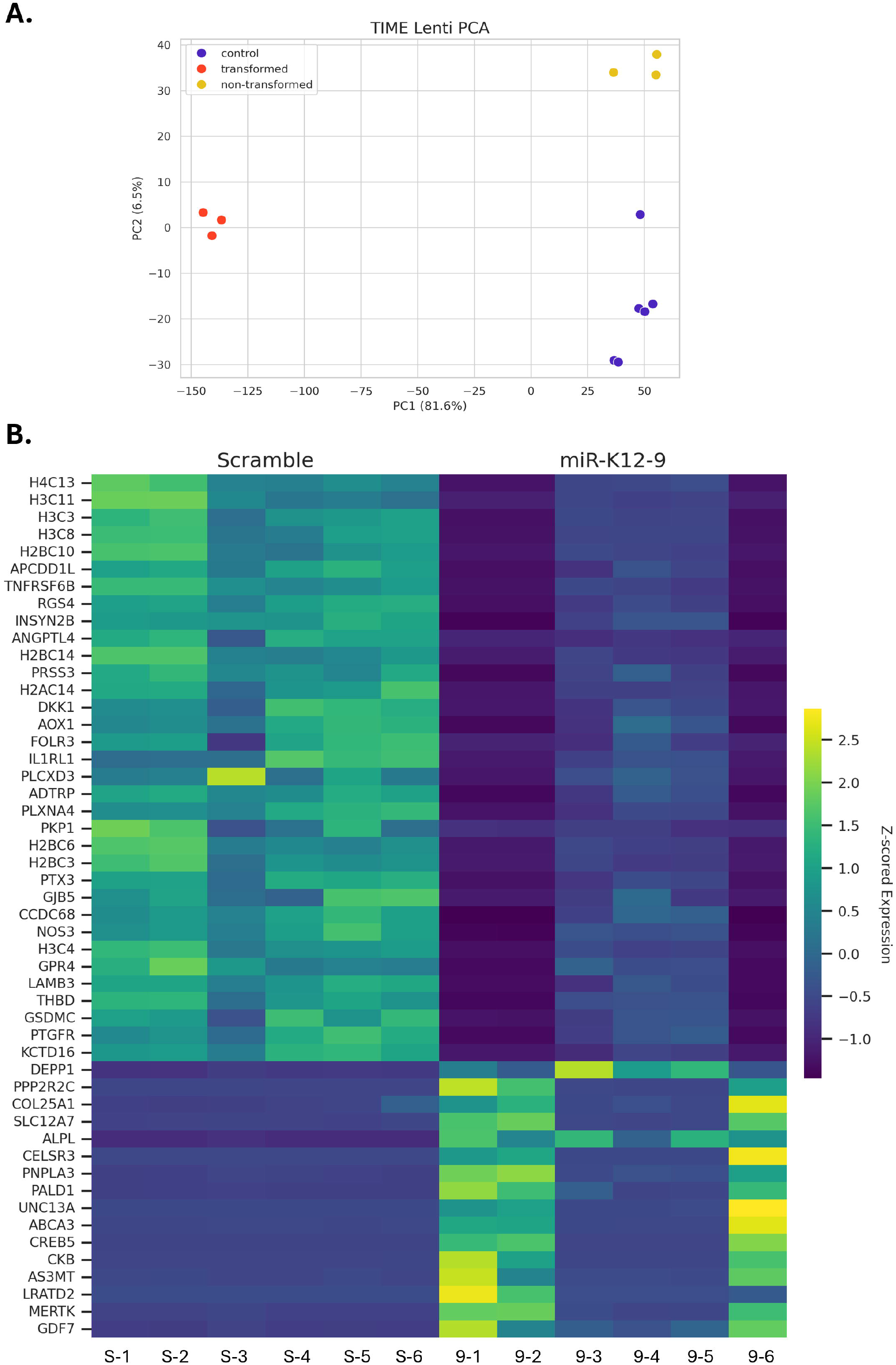
Results of assays characterizing the control miRNA- and miR-K12-9-expressing TIME cells. A. Expression of mature miR-K12-9-5p was quantified by qPCR. The data are shown as average fold change over WT KSHV-infected cells. Error bars represent standard deviation. The experiment was performed with RNA from 3 separate preparations. B. Cells were seeded in 96-well plates and incubated for 72h, at which point CellTiter Glo reagents (Promega) were added and luminescence was quantified. The data are displayed as relative luminescence. Error bars represent standard deviation. C. Cells were suspended in diluted agar and distributed to 6-well plates. After covering with cell culture medium, the cells were left to incubate for 2 weeks. Each well was photographed and colonies were counted using ImageJ. D and E. NOD/SCID mice were injected in the left flank with control miRNA-expressing cells and in the right flank with miR-K12-9-expressing cells. Three mice were used per biological replicate. Only mice injected with miR-K12-9 BRs 1, 2, and 6 developed tumors. D. Graph showing the growth of tumors over 6 weeks. E. Images of excised tumors.

This pattern was further supported by hierarchical clustering of differentially expressed genes (Figure 5B), which demonstrated that miR-K12-9 expression alone induces a coordinated transcriptional program that is partially shared across all expressing cell lines. Importantly, this program was markedly amplified in transformed cells, which exhibited more extreme expression changes across many of the same gene sets. Together, these findings indicate that miR-K12-9 expression establishes a distinct transcriptional state that is further reinforced during oncogenic transformation. Rather than representing entirely separate programs, the transcriptional profiles of non-transformed and transformed cells appear to lie along a continuum, suggesting that miR-K12-9 establishes a permissive transcriptional landscape for oncogenic transformation, with additional factors driving progression to a fully transformed state.

Although these findings support a role for miR-K12-9 in promoting oncogenic transformation, the TIME cell model introduces an important caveat, as these cells are telomerase-immortalized and may already possess features that facilitate transformation. This raised the possibility that miR-K12-9 acts in cooperation with pre-existing cellular alterations, consistent with a “two-hit” model in which telomerase expression provides a permissive background. To determine whether miR-K12-9 alone is sufficient to drive transformation in a less primed system, we repeated these experiments in primary Human Umbilical Vein Endothelial Cells (HUVECs).

HUVECs were transduced with miR-K12-9 or control lentiviruses under the same conditions used for TIME cells. Notably, a subset of miR-K12-9-expressing HUVEC cultures developed a rapidly proliferating phenotype within several weeks, similar to that observed in TIME cells. This phenotype emerged in approximately 50% of miR-K12-9-transduced wells, whereas control-transduced cells maintained normal growth characteristics (Figure 6A, 6B). As expected, expression of mature miR-K12-9-5p and -3p was detected in all transduced cells, regardless of phenotype. However, only cells exhibiting the fast-growing phenotype demonstrated anchorage-independent growth in soft agar, consistent with acquisition of transformative properties (Figure 7A, 7B).

**Figure 6.**
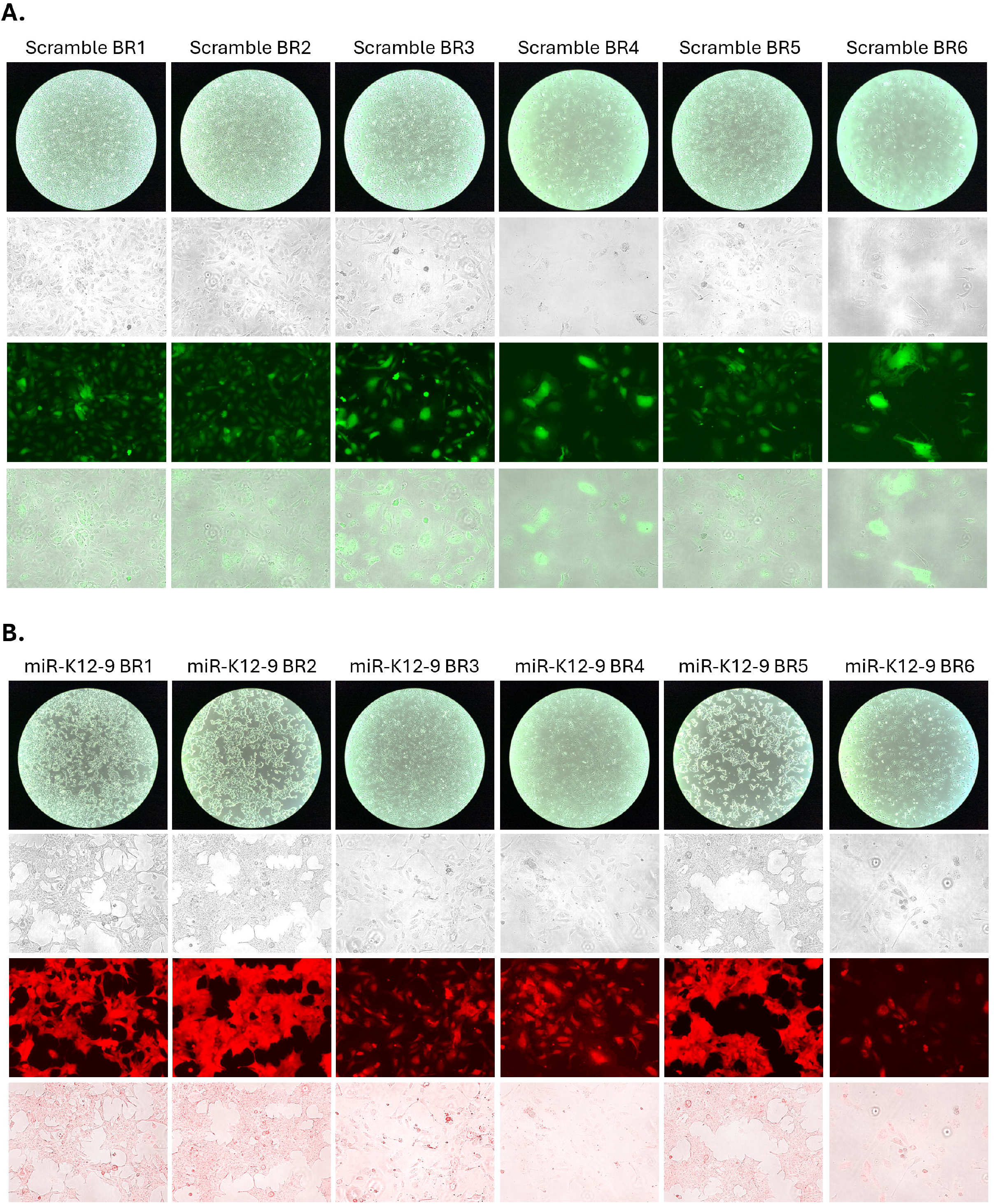
Heatmap showing the top differentially expressed genes between control miRNA-and miR-K12-9-expressing cells. Expression was normalized by Z-score.

**Figure 7.**
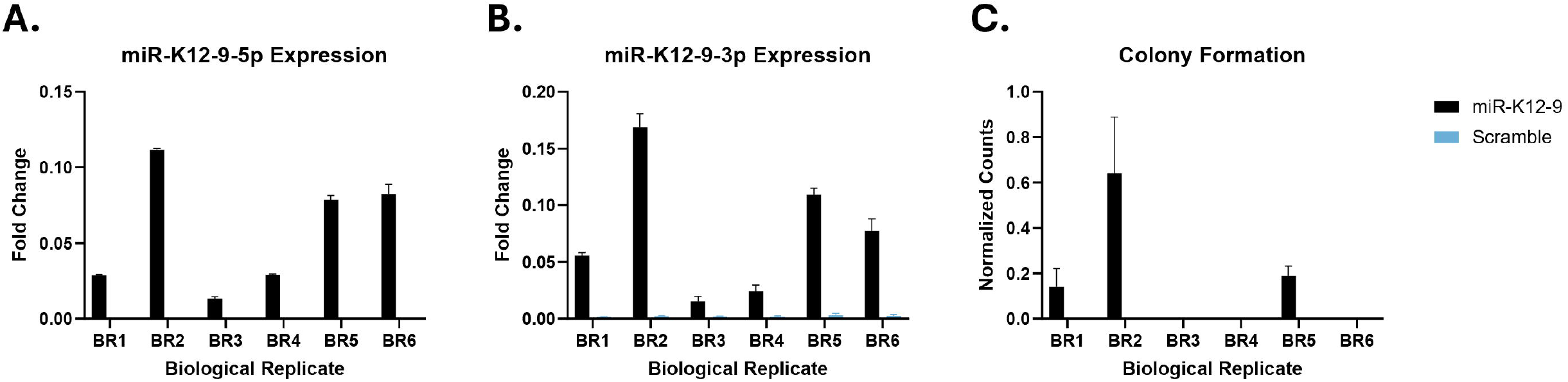
Images of HUVEC cells transduced with control (GFP) and miR-K12-9 (mCherry) lentiviruses. HUVEC cells in 6-well plates were transduced in triplicate with either control miRNA- or miR-K12-9-expressing lentivirus. Each transduced well is considered a biological replicate (BR). The photographs shown were taken 27 days post-transduction, after puromycin selection and expansion into 10cm plates.

**Figure 8.** Results of assays characterizing the control miRNA- and miR-K12-9-expressing HUVEC cells. A. Expression of mature miR-K12-9-5p was quantified by qPCR. The data are shown as average fold change over WT KSHV-infected cells. Error bars represent standard deviation. The experiment was performed with RNA from 3 separate preparations. B. Cells were suspended in diluted agar and distributed to 6-well plates. After covering with cell culture medium, the cells were left to incubate for 2 weeks. Each well was photographed and colonies were counted using ImageJ.

The graded transcriptional changes observed across miR-K12-9-expressing cells, culminating in transformation in a subset of replicates, suggested that additional factors may cooperate with miR-K12-9 to drive full oncogenic conversion. One potential explanation is that lentiviral integration events could influence the expression of nearby oncogenes or tumor suppressors (32). To address this possibility, integration sites were mapped in the original miR-K12-9-expressing cell lines. The transformed line BR1 contained three integration sites within introns of *GRB2, SETD2*, and *TSPAN7*, while the transformed line BR2 harbored a single integration within an intron of *RYK*. However, RNA-seq analysis revealed no significant changes in expression of these genes relative to control cells. Furthermore, read coverage upstream and downstream of each integration site was comparable between samples, indicating no detectable disruption of transcript structure or splicing (Fig. S9). The control and non-transformed cells had many integration sites, each with low numbers, suggesting that these cell lines are not clonal. Together, these findings suggest that lentiviral integration is unlikely to account for the observed transformation phenotype, supporting a model in which miR-K12-9 establishes a permissive transcriptional state that requires additional, non-integration-related factors to drive full transformation.

Overall, these findings indicate that miR-K12-9 expression can drive key features of cellular transformation even in primary endothelial cells, independent of prior immortalization. At the same time, the incomplete penetrance of the phenotype suggests that additional cellular factors or stochastic events contribute to the transition to a fully transformed state. Together, these results further support a model in which miR-K12-9 strongly biases cells toward oncogenic transformation while requiring cooperating conditions for full phenotypic conversion.

## Discussion

In this study, we present RNA-seq and small RNA-seq datasets derived from a complete panel of KSHV miRNA knockout mutants in endothelial cells. These datasets are intended to serve as a resource for the broader KSHV research community, complementing prior phenotypic analyses and enabling further exploration of miRNA-mediated regulation during KSHV infection. Consistent with earlier observations from our laboratory, the combined transcriptional and phenotypic data indicate that KSHV miRNAs exhibit both redundancy and specialization, collectively influencing a wide range of cellular pathways while maintaining distinct individual functions.

Among the viral miRNAs examined, miR-K12-9 emerged as particularly unusual. In contrast to the smaller effects observed for other miRNA knockouts, loss of miR-K12-9 resulted in widespread transcriptional disruption, affecting thousands of genes and multiple major biological pathways. This broad and multifaceted impact, together with the pronounced phenotypic changes observed previously, prompted further investigation into the functional role of this miRNA.

Unexpectedly, sustained ectopic expression of miR-K12-9 in endothelial cells was sufficient to drive cellular transformation in a subset of cases. This was observed across multiple independent biological replicates in both immortalized TIME cells and primary HUVECs, showing that the effect does not rely on prior immortalization. Transformed cells exhibited hallmark features of oncogenesis, including increased proliferation, anchorage-independent growth, and tumor formation *in vivo*.

Transcriptomic analysis of miR-K12-9-expressing cells provides insight into this phenomenon. RNA-seq revealed that miR-K12-9 induces a coordinated transcriptional program that is detectable even in non-transformed cells and is further amplified in transformed populations. Rather than representing discrete states, the transcriptional profiles of control, non-transformed, and transformed cells appear to lie along a continuum, consistent with a model in which miR-K12-9 establishes a permissive transcriptional landscape that predisposes cells toward transformation. The transition to a fully transformed phenotype likely depends on additional cooperating events.

One possible cofactor for transformation could be insertional mutagenesis caused by lentivirus integration. However, analysis of integration sites in transformed cell lines did not reveal changes in expression or transcript structure of the genes containing the integration sites or nearby genes, arguing against a primary role for insertional mutagenesis. Instead, the observed heterogeneity in transformation may reflect intrinsic differences within the starting cell populations. TIME cells, while derived from a single donor, are reported to develop karyotypic instability during extended passage (33) and the HUVECs used in this study were obtained from pooled donors. It is therefore possible that certain genetic or epigenetic backgrounds are more permissive to miR-K12-9-driven transformation than others. Alternatively, stochastic factors or differences in cellular state at the time of transduction may contribute to the observed variability.

Several important questions remain to be addressed. First, it is unclear whether continued expression of miR-K12-9 is required to maintain the transformed phenotype, or whether it functions primarily as an initiating factor. Future studies using a tet-off expression system will be necessary to determine whether transformation is reversible upon loss of miR-K12-9. Second, while the present work focuses on vascular endothelial cells, it will be important to assess whether similar effects can be observed in other relevant cell types, particularly lymphatic endothelial cells, which are the most likely origin of KS spindle cells (34). Finally, the identification of cooperating factors that enable full transformation represents a critical next step. Modulating specific cellular pathways in non-transformed miR-K12-9-expressing cells may help define the conditions required for oncogenic conversion. Additionally, the non-transformed cells present an opportunity to test other viral gene products for the ability to cooperatively drive oncogenesis.

In summary, our findings identify miR-K12-9 as a potent regulator of host transcriptional programs and demonstrate that, in the absence of other viral gene products, it is capable of driving oncogenic transformation in endothelial cells under permissive conditions. To our knowledge, this represents the first report of a KSHV miRNA with the capacity to induce transformation of endothelial cells, a relevant model for KS. These results expand the known functional repertoire of viral miRNAs and highlight miR-K12-9 as a key factor in KSHV-associated oncogenesis.

## Materials and Methods

### Cell Culture

Telomerase-Immortalized Microvascular Endothelial (TIME) cells, a generous gift from Dr. Michael Lagunoff, were grown in Endothelial Cell Growth Medium-2 (EGM-2) MV with supplements provided in the BulletKit (Lonza, # CC-3202). Lenti-X 293T cells (Takara Bio, 632180) were grown in DMEM with 10% tetracycline-free FBS and 1% penicillin/streptomycin. The cell culture medium for HUVECs (Lonza, #CC-2519) was prepared from the EGM Endothelial Cell Growth Medium BulletKit (Lonza, #CC-3124). An additional 25 ml of FBS was added for a final FBS concentration of 7%.

### Establishment of miRNA and inhibitor expressing cell lines

Plasmids encoding the desired transgenes were ordered from Vector Builder. These were pLV-miR-K12-9, containing the genomic hairpin sequence of miR-K12-9 surrounded by 100bp on either side; pLV-Scramble, containing a non-targeting shRNA; pLV-Ind-miR-K12-9-5p-AS, encoding an inducible miR-K12-9-5p inhibitor; and pLV-Ind-Scramble, encoding an inducible non-targeting shRNA. Lentiviruses containing the transgenes were created with the Lenti-X system from Takara Bio. Lenti-X 293T cells were transfected with the lentiviral vectors using Lenti-X Packaging Single Shots according to the manufacturer’s instructions. Supernatants were collected after 48hr and cleared by sedimentation. Two milliliters of cleared supernatant were transferred to each well of a 6-well plate seeded the day before at a density of 1 × 10^5^ cells/ml. The plates were centrifuged at 2500rpm for 1.5hr at 30°C and then returned to 37°C o/n. The next day the supernatant was replaced with fresh medium. Seventy-two hours post-transduction puromycin was added to the cell culture medium at a concentration of 1ng/µl. Transduced cells were expanded and maintained at the same concentration of puromycin.

### RNA Isolation and DNase Treatment

Cells from 80-100% confluent 10cm plates were resuspended in 1ml of RNA STAT-60 (Amsbio, #CS-502) and transferred to 1.5ml tubes. Samples were either stored at -80°C or processed directly. RNA was prepared according to the RNA STAT-60 manufacturer’s protocol. The final RNA pellets were resuspended in 20-50µl of DEPC water and incubated at 55°C for 10min. Concentrations were measured by NanoDrop and RNA was diluted to 1µg/µl.

DNase treatment was carried out in one of two ways. For RNA destined for qPCR, 10µg of RNA was combined with 5µl of DNase I (NEB), 2µl of DNaseI Buffer, and DEPC-treated water up to 20µl. Samples were incubated at 37°C for 30min. The DNase reaction was terminated by adding 2µl of 25mM EDTA to each tube followed by incubation at 65°C for 10min. DNase-treated RNA was either polyadenylated or directly used in reverse transcription followed by qPCR. RNA intended for RNA-seq was not incubated with EDTA but rather re-extracted. The volume in each tube was brought up to 250µl with DEPC water, at which point 250µl of Phenol/Chloroform/Isoamyl Alcohol (25:24:1) (ThermoFisher, #AM9730) was added and the samples were shaken for 8min at 1400rpm and 25°C. This was followed by centrifugation for 10min at 18,000 x g. The aqueous layers were transferred to new tubes to which 500µl Ethanol/Isopropanol 1:1, 20µl 3M sodium acetate, and 2µl GlycoBlue (ThermoFisher, #AM9516) were added. Tubes were inverted several times to mix and stored at -20°C o/n. The next day, samples were centrifuged for 30min at 21,000 x g and 4°C. Pellets were washed once with 900 µl of ice-cold 80% ethanol, centrifuged 10min at 18,000 x g and 4°C, and allowed to air-dry for 5-10min. RNA pellets were resuspended in 20µl of DEPC water. Concentrations were measured by NanoDrop and RNA was diluted to 1µg/µl in DEPC water and stored at -80°C.

### Polyadenylation, Reverse Transcription, and qPCR

Polyadenylation of DNase-treated RNA intended for miRNA qPCR was performed with a Poly(A) Polymerase Tailing Kit (Biosearch Technologies, # PAP5104H) starting with 2µg of DNase-treated RNA. RNA was reverse-transcribed using SuperScript IV (Invitrogen) with the miRNA oligo(dT) primer (see Table X for all primer sequences) for poly(A) RNA and random hexamers for non-poly(A) RNA.

All qPCR was performed on a Roche LightCycler 96 using DNA Green Essential Master Mix (Roche). For quantification of coding genes of interest, the following qPCR program was used: preincubation at 95°C for 30s, 45 cycles of 95°C for 20s, 56°C for 20s, and 72°C for 30s, followed by a melting curve. For miRNA qPCR the program used was: 95°C for 1200s, 40 cycles of 95°C for 15s and 60°C for 60s, and a final melting curve. Results were analyzed using the LightCycler 96 software version 1.1.

### Colony Formation Assays

Colony formation assays were performed as in Du et al with several modifications (35). Freshly prepared 5% LB-agar in PBS was placed in a 50° C water bath while still liquid and allowed to equilibrate for at least 1h. At that time 4 ml of 5% agar was combined with 36 ml of EGM-2 MV pre-warmed to 37° C. One milliliter of this mixture was distributed to each well of 6 6-well plates and the agar was allowed to solidify at room temperature for at least 30 min. Meanwhile, TIME cells were trypsinized and resuspended at a concentration of 1 × 10^3^ cells/ml in EGM-2 MV. Three-hundred microliters of 5% agar were combined with 4.7 ml of cell suspension and 1 ml of this mixture was dispensed to each of 3 wells on top of the previous agar layer. The new layer was allowed to solidify at RT for 30 min before 1 ml of EGM-2 was added to each well and the plates were returned to 37° C. Colonies were photographed after 3 weeks and counted using ImageJ.

### Cell Proliferation Assays

Cells were seeded at a density of 1 × 10^4^ cells/ml in opaque-walled 96-well plates. Proliferation was measured after 72 h using the CellTiter-Glo Luminescent Cell Viability Assay (Promega, #G7570). Each well received 20 µl of CellTiter-Glo reagent and measurements were performed on a FluoStar Optima instrument (BMG Labtech).

### Mouse Tumorigenesis Assays

Scramble shRNA or miR-K12-9-expressing TIME cells were resuspended in PBS at a density of 1 × 10^7^ cells/mL. One hundred microliters of the cell suspension were combined 1:1 with growth factor reduced Matrigel (Corning?). NOD/SCID mice were injected subcutaneously into the flank with 100 µl of the Matrigel mixture. The left flanks received the scramble shRNA-expressing cells while the right flanks received the miR-K12-9-expressing cells.

Mice were palpated daily for tumors. Any tumors that appeared were measured on a weekly basis and the volume was calculated. Mice were sacrificed after 6 weeks or when the tumor volume reached 1500 cubic millimeters. Excised tumors were divided into 3 sections. One section was placed in formalin and later used to prepare hematoxylin and eosin-stained slides. The other sections were either flash-frozen in liquid nitrogen or used fresh in scRNA-seq.

### RNA-seq

Starting with 1 µg of DNase I-treated RNA with a RIN of at least 7, ribosomal RNA was removed using the NEBNext® rRNA Depletion Kit (Human/Mouse/Rat) (NEB, # E6310). Libraries for high-throughput sequencing were prepared with the NEBNext® Ultra II Directional RNA Library Prep Kit for Illumina (NEB, #E7760). Sequencing was performed on an Illumina NovaSeq6000 at the University of Florida Interdisciplinary Center for Biomedical Research (UF ICBR).

### Single-Cell RNA-seq

Freshly resected mouse tumors were minced into approximately 1 mm pieces and placed in 5 ml of digest solution (10 mM HEPES, 5mM CaCl_2_, 5% FBS, 50 U/ml DNase I, 1.5 mg/ml collagenase A in Hank’s balanced salt solution) pre-warmed to 37°C. The tissue pieces were incubated at 37°C while rotating and vortexed every 5 min. The solution was added to 25 ml of PBS, vortexed for a further 30 s, and passed through a 70 µm screen into a new tube. Cells were pelleted and resuspended in 2 ml of ACK lysis buffer for 3 min, at which point 25 ml of PBS was added and cells were pelleted again. Finally, the cells were resuspended in 10 ml of complete EGM-2 MV and transferred to a 10 cm plate. The cultures were allowed to grow to confluency prior to fixation. Cells were fixed using the Evercode Cell or Nuclei Fixation kit v3 (Evercode, ECFC3300) and stored at -80°C until library preparation. Libraries were prepared with the Parse Evercode WT kit v3 following the manufacturer’s instructions and sequenced on a NovaSeq6000 at UF ICBR.

### Bioinformatics Analysis

RNA-seq and small RNA-seq data were analyzed at the Tulane Cancer Center Next Generation Sequencing Analysis core. Raw fastq files were quality checked with FastQC v0.11.7 (36). Small RNA-seq data was analyzed with miRExpress (37). For regular RNA-seq, reads were aligned against the hg38 reference genome assembly GRCh38.p14 (RefSeq GCF_000001405.40) plus the GK18 KSHV genome (RefSeq NC_009333) using STAR aligner v2.5.2a (38). Differential expression analysis was performed with DESeq2 and Sleuth while splicing analysis was carried out using rMATS v4.1.1 (39,40,41). Finally, gene set enrichment analysis was completed with GSEA v4.3.2 (42).

Single-cell RNA-seq data was first analyzed with the Parse Trailmaker software. Filtered matrices were further analyzed with scanpy to generate unsupervised Leiden clusters and perform differential expression analysis (43). Gene set enrichment analysis was performed with gseapy (44). Graphs were made with matplotlib (45).

For integration site sequencing, reads containing the LTR sequence were extracted and the LTR and Illumina read 1 adapter sequences were trimmed with Cutadapt (46). The trimmed reads were aligned to the lentiviral vector sequence with Bowtie2 and only reads that failed to align were kept (47). These reads were then aligned to hg38 with BWA, using the hg38 assembly specified by this program (48). Bigwig files were generated with samtools and visualized on IGV (49, 50). For pinpointing integration sites, bedtools was used to merge nearby sites and output bed files (51).

Except for bulk RNA-seq analysis, custom scripts for implementing command line tools were created using ChatGPT and can be found on GitHub.

### Integration Site Identification

Genomic DNA was extracted from cells with the DNeasy Blood & Tissue Kit (Qiagen) and digested with DNA Fragmentase (NEB) to yield fragments with an average size of approx. 200bp. Sequencing libraries were prepared with the NEBNext Ultra II DNA Library Prep Kit for Illumina (NEB) following the LM-PCR strategy from Kim et al (52). Briefly, the NEBNext kit reagents were used for end repair and adapter ligation. This was followed by 2-step nested PCR with primers specific for the LTR and Illumina adapter. Steps 1 and 2 had the same program: 98°C for 30s, *98°C for 10s, 65°C for 30s, 72°C for 30s, repeat from * 25x, and 72°C for 2 min. Primers from the NEBNext Multiplex Oligos for Illumina Set 2 were used in a final PCR following the manufacturer’s program with 30 cycles. Libraries were sequenced on a NovaSeqX 6000 with 150bp paired-end sequencing.

## Figure Legends

Figure S1. Volcano plots showing differentially expressed genes between miRNA KO mutants miR-K12-1 – miR-K12-6 vs. WT.

Figure S2. Volcano plots showing differentially expressed genes between miRNA KO mutants miR-K12-7 – miR-K12-12 vs. WT.

Figure S3. Bubble plots showing gene set enrichment analysis results for miRNA KO mutants miR-K12-1 – miR-K12-6.

Figure S4. Bubble plots showing gene set enrichment analysis results for miRNA KO mutants miR-K12-7 – miR-K12-12.

Figure S5. Heatmaps showing top 50 differentially expressed genes for miRNA KO mutants miR-K12-1 – miR-K12-4 vs WT.

Figure S6. Heatmaps showing top 50 differentially expressed genes for miRNA KO mutants miR-K12-5– miR-K12-8 vs WT.

Figure S7. Heatmaps showing top 50 differentially expressed genes for miRNA KO mutants miR-K12-9 – miR-K12-12 vs WT.

Figure S8. Integrated Genomics Viewer (IGV) tracks showing RNA-seq read coverage of genes hosting lentivirus integration sites. Integration sites are marked with a black vertical bar below the corresponding track. A. GRB2, B. SETD2, C. TSPAN7, D. RYK.

