## Supplementary figures and images for "KSHV miR-K12-9 Induces Transformation of Immortalized and Primary Endothelial Cells"

### Figure S1

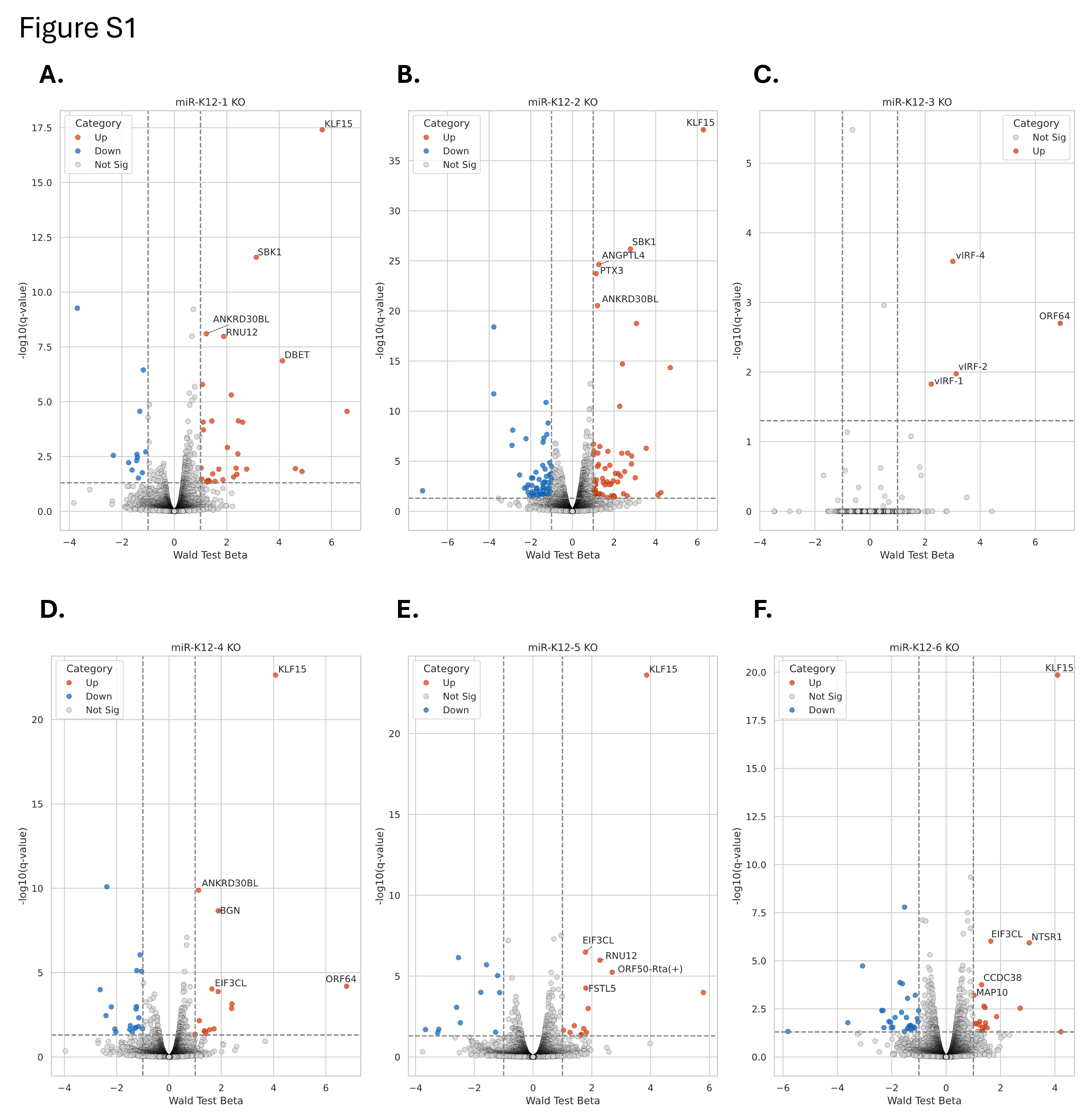

### Figure S2

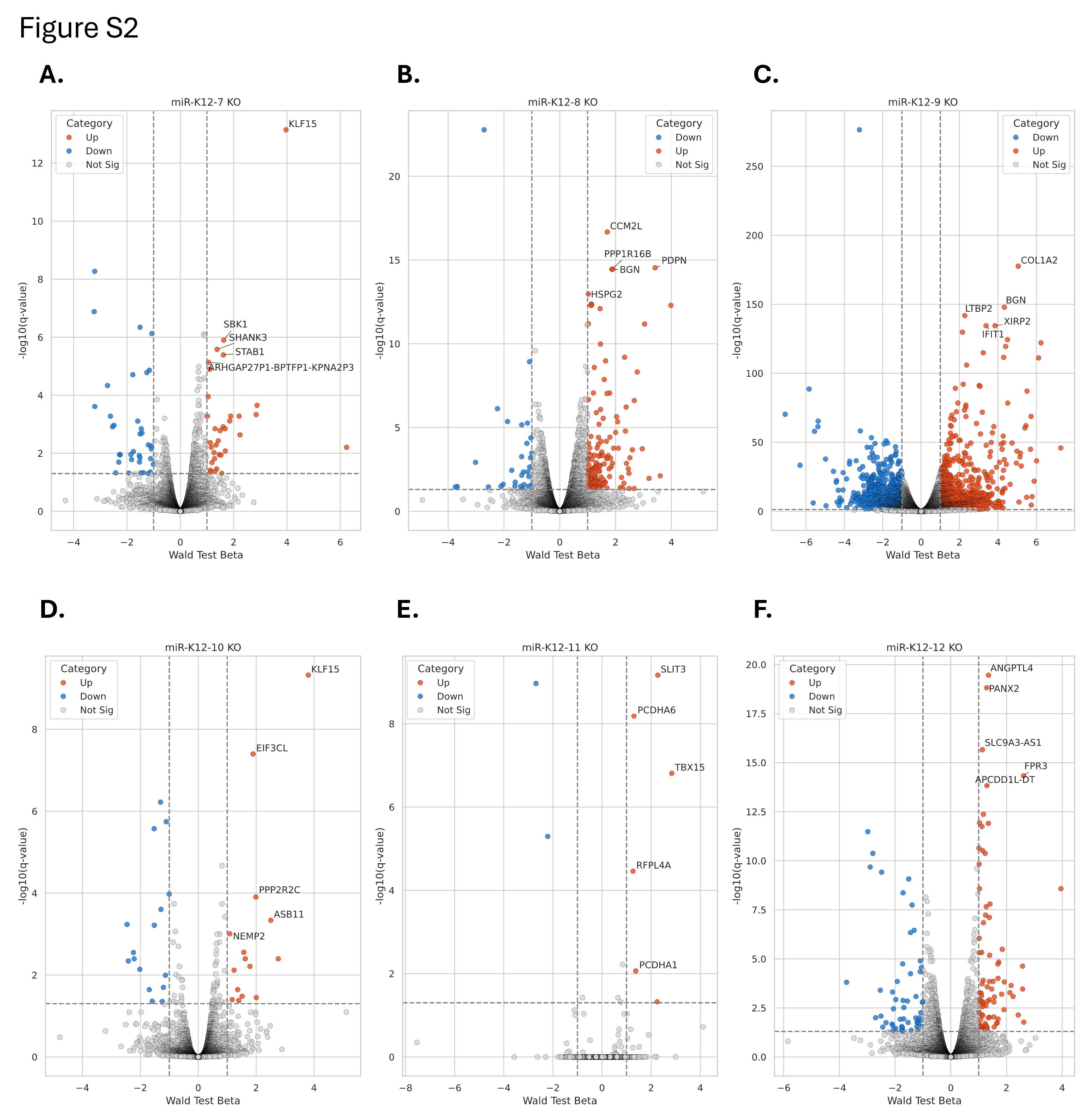

### Figure S3

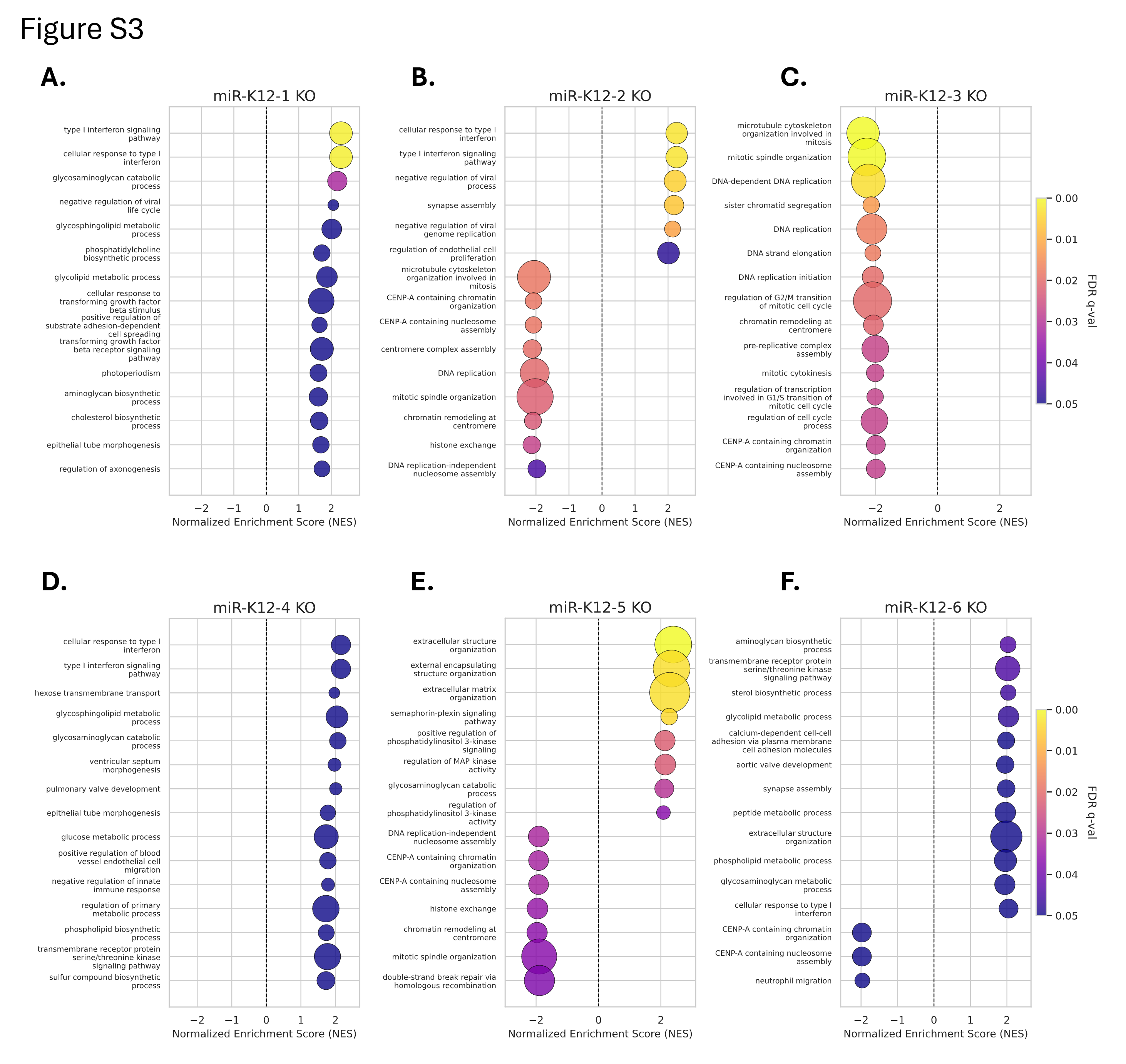

### Figure S4

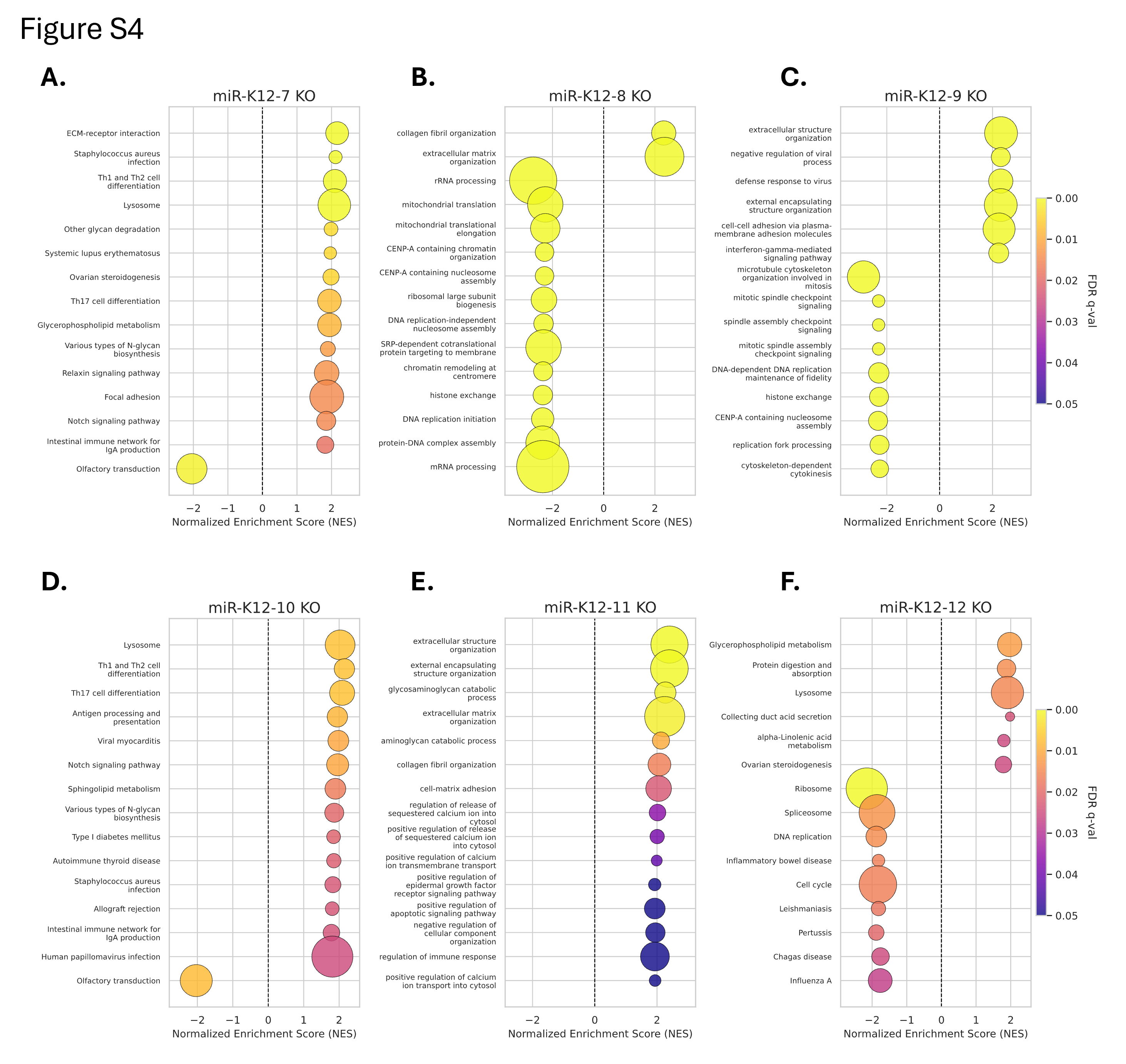

### Figure S5

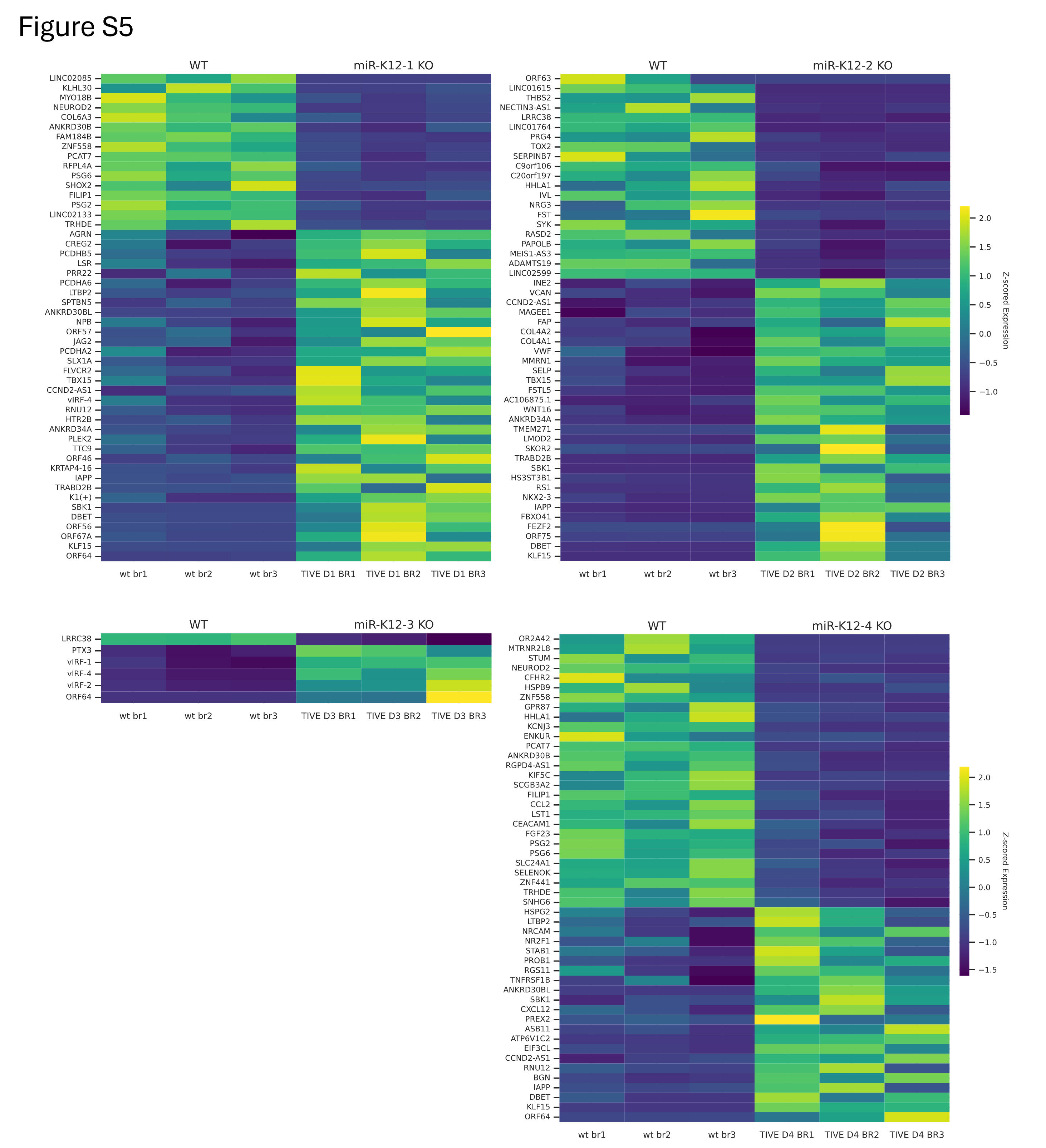

### Figure S6

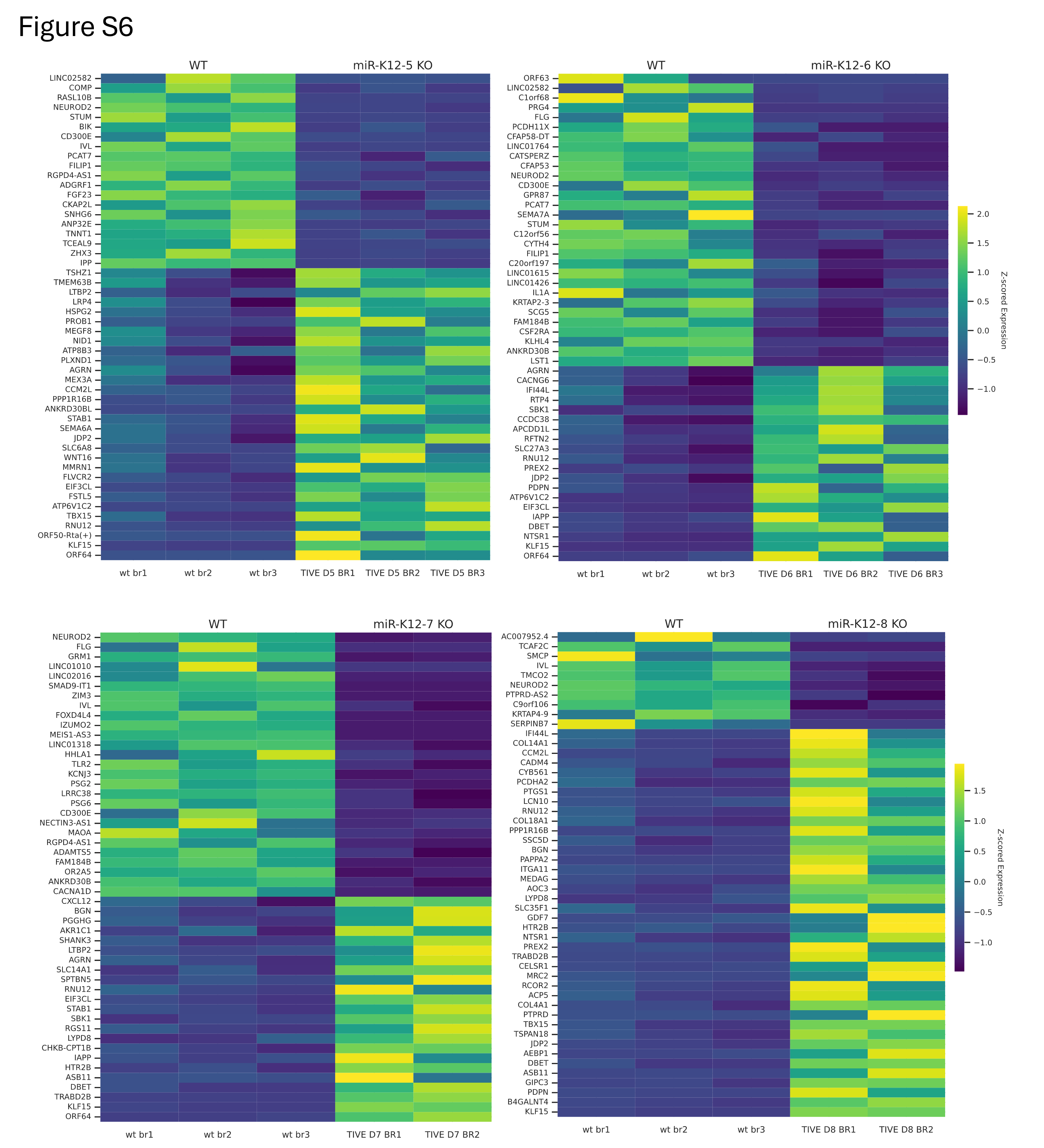

### Figure S7

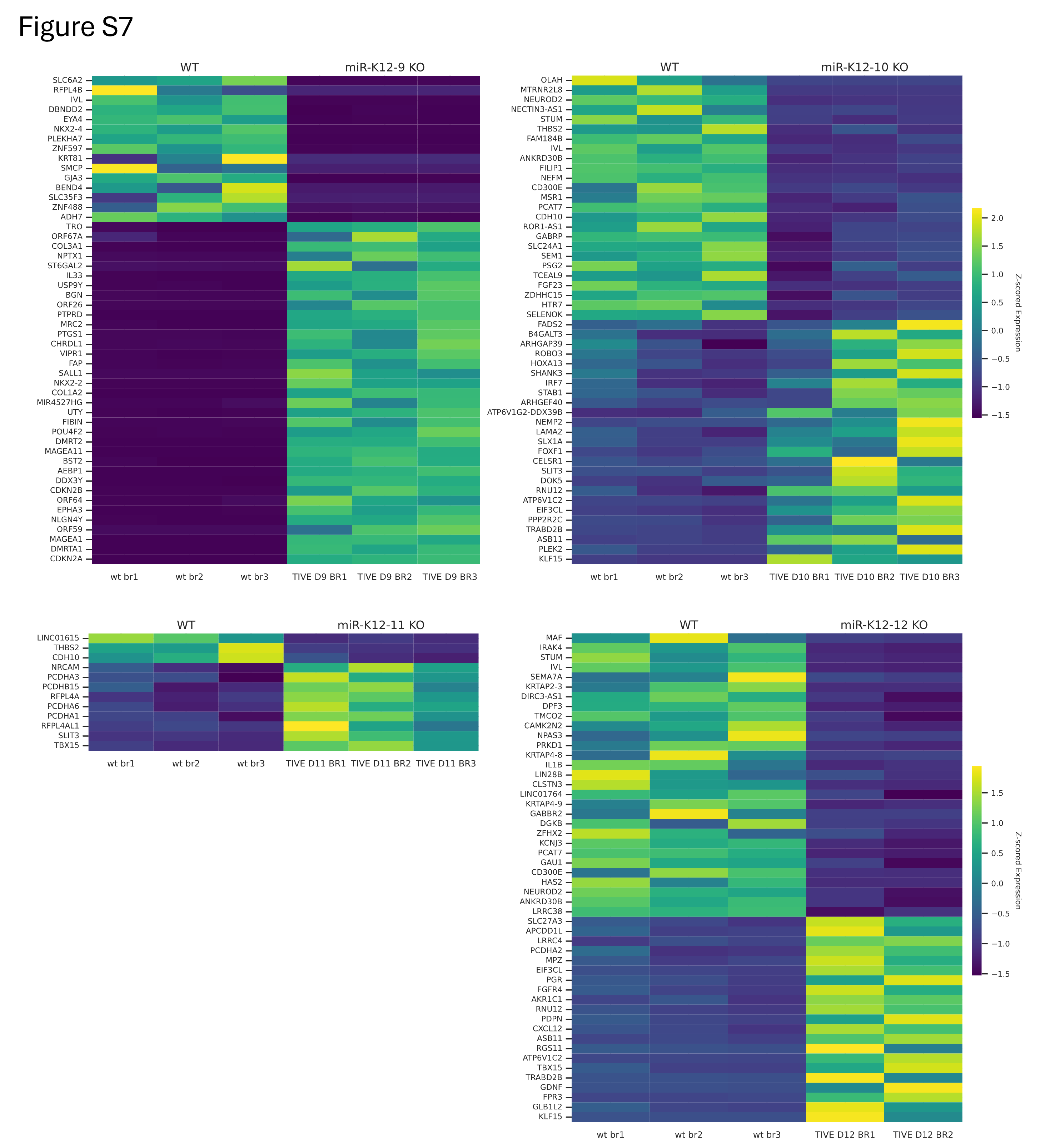

### Figure S8

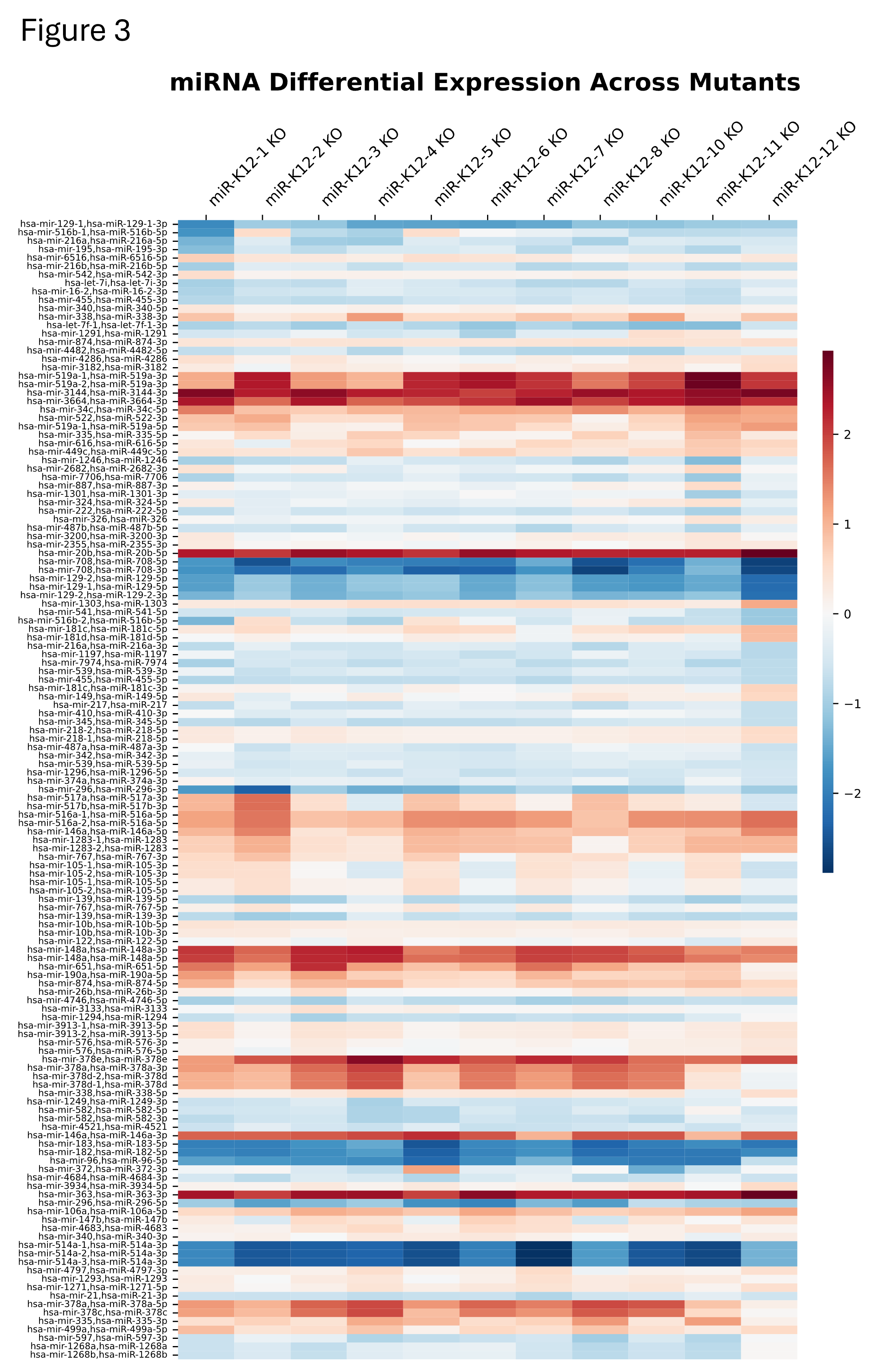

### Figure S9

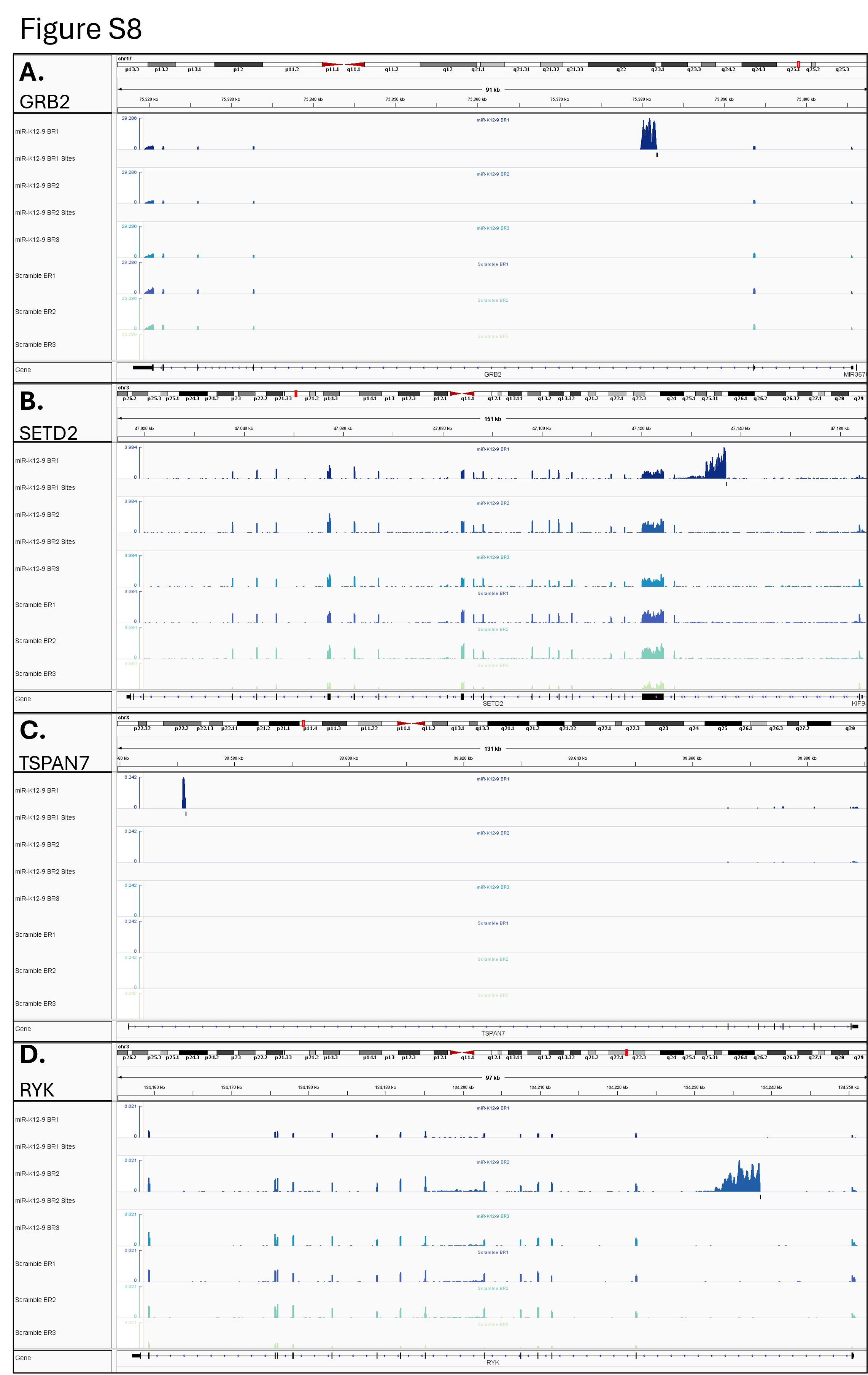
